# Pannexin-1 Channels govern the Generation of Necroptotic small Extracellular Vesicles

**DOI:** 10.1101/537753

**Authors:** Tiphaine Douanne, Gwennan André-Grégoire, Magalie Feyeux, Philippe Hulin, Julie Gavard, Nicolas Bidère

## Abstract

The activation of mixed lineage kinase-like (MLKL) by receptor-interacting protein kinase-3 (RIPK3) controls the execution of necroptosis, a regulated form of necrosis that occurs in apoptosis-deficient conditions. Active oligomerized MLKL triggers the exposure of phosphatidylserine residues on the cell surface and disrupts the plasma membrane integrity by forming lytic pores. MLKL also governs the biogenesis and shedding of proinflammatory small extracellular vesicles (EVs) during the early steps of necroptosis, however the molecular basis is unknown. Here, we find that MLKL oligomers activate plasma membrane Pannexin-1 (PANX1) channels, concomitantly to the loss of phosphatidylserine asymmetry. This plasma membrane “leakiness” requires the Rab GTPase Rab27 isoforms, which usher the small EVs to their release. Conversely, PANX1 knockdown disorganized the small EVs machinery and precludes vesicles extrusion. These data identify a novel signaling nexus between MLKL, Rab27, and PANX1, and propose ways to interfere with small EV generation.

## INTRODUCTION

Necroptosis is a programmed lytic cell death pathway that occurs in apoptosis-deficient conditions [1]. In contrast to apoptosis, which is immunologically silent as cellular content remains impermeable and is rapidly engulfed by surrounding phagocytic cells, necroptotic cells produce proinflammatory cytokines and release cytosolic proinflammatory material in the extracellular milieu as a consequence of the loss of plasma membrane (PM) integrity [2]. Although the engagement of several immunoreceptors culminate in necroptosis, this form of death has been best studied upon ligation of the death receptor family member TNF receptor 1 (TNFR1) [2]. When caspases are blocked, TNFR1 signaling assembles a filamentous death-inducing complex of the Ser/Thr kinases RIPK1, RIPK3 and the pseudokinase MLKL coined necrosome, which culminates in the phosphorylation of MLKL by RIPK3 [3–9]. This causes MLKL to adopt a conformational change required for its multimerization [10]. The binding of negatively charged phospholipids such as high-order inositol phosphates to oligomerized MLKL allows the pseudokinase to insert into the PM and compromise cell-membrane integrity leading to the cell’s demise [10–14].

Shortly before PM collapses, MLKL oligomers trigger the exposure of phosphatidylserine (PS) residues at the cell surface, acting as “eat-me” signals for the surrounding immune cells [15,16]. However, cells with phosphorylated MLKL and flipped PS are not necessarily committed to death and can “resuscitate” provided the stimulus is removed in a timely manner [16,17]. MLKL also governs the dynamic biogenesis and release of small extracellular vesicles (EVs) during the early stages of necroptosis by cells which are otherwise viable [15,18]. These membrane vesicles containing cytosolic material emerge as potent vectors of intercellular communication [19]. How MLKL controls these many functions continues to be elucidated. Hints may come from the discovery that active MLKL targets intracellular organelles in addition to the PM [3], and was demonstrated to regulate constitutive endosomal trafficking [18]. Furthermore, MLKL silencing causes a drastic reduction in the number of nascent intracellular small EVs, also called intraluminal vesicles (ILVs), which are no longer expelled into the extracellular space [18].

The mechanism by which MLKL licenses the biogenesis and the release of small EVs out of the cells remains poorly understood. Here, we find that MLKL oligomers also promote the activation of PANX1 hemichannels via the Rab27 small GTPase subfamily during the early stages of necroptosis. Although this Rab27-PANX1 axis appears dispensable for the proper execution of the necroptotic program, it tightly regulates the generation of small EVs.

## RESULTS

### MLKL promotes “Leakiness” of the Plasma Membrane during Necroptosis

HT-29 colon cancer cells, which constitute a classical model for necroptosis [4,5] were treated with TNF*α* (T) in the presence of the pan-caspase inhibitor Q-VD-OPh (Q), together with the Smac mimetic Birinapant (S) [20]. Necroptosis was then visualized by depletion of intracellular ATP, crystal violet staining, staining of PS residues on the outer face of PM with Annexin-V (A5), and uptake of the vital cell-impermeant dye TO-PRO-3 [4,5,15–17]. As expected [3,6], cell death was efficiently prevented by either the RIPK1 inhibitor Nec-1s or by silencing MLKL (Fig. 1A-D). Supporting an initial observation by Gong *et al* [17], staining with TO-PRO-3 discriminated a population of “leaky” cells with a dim staining from dead cells with a high incorporation of the dye (Fig. 1C and 1E). Of note, those TO-PRO-3^Dim^cells were mostly A5-positive (Fig. 1C). Moreover, “leakiness” preceded full permeabilization to the dye seen after 24 hours, which likely reflects the loss of PM integrity (Ref. [17], Fig. 1F and 1G). Importantly, the early uptake of TO-PRO-3 was selective, as it was not accompanied by a staining with propidium iodide (PI), although of smaller molecular weight (Fig. 1H and 1I). Further militating against a general rupture of the PM, cytosolic lactate deshydrogenase (LDH) was only modestly released in the extracellular milieu during early steps of necroptosis (Fig. 1J). Time-lapse microscopy to monitor A5 staining and TO-PRO-3 uptake further unveiled that kinetics of PS exposure and TO-PRO-3 uptake during necroptosis slightly differed (Fig. 1K, and Movies S1 and S2). Although both events started 2 hours post-stimulation, PS exposure occurred suddenly as compared to the gradual uptake of TO-PRO-3. Altogether, this suggests that the PM of cells becomes “leaky” to TO-PRO-3 in an MLKL-dependent manner during the early onset of necroptosis.

**Figure. 1.**
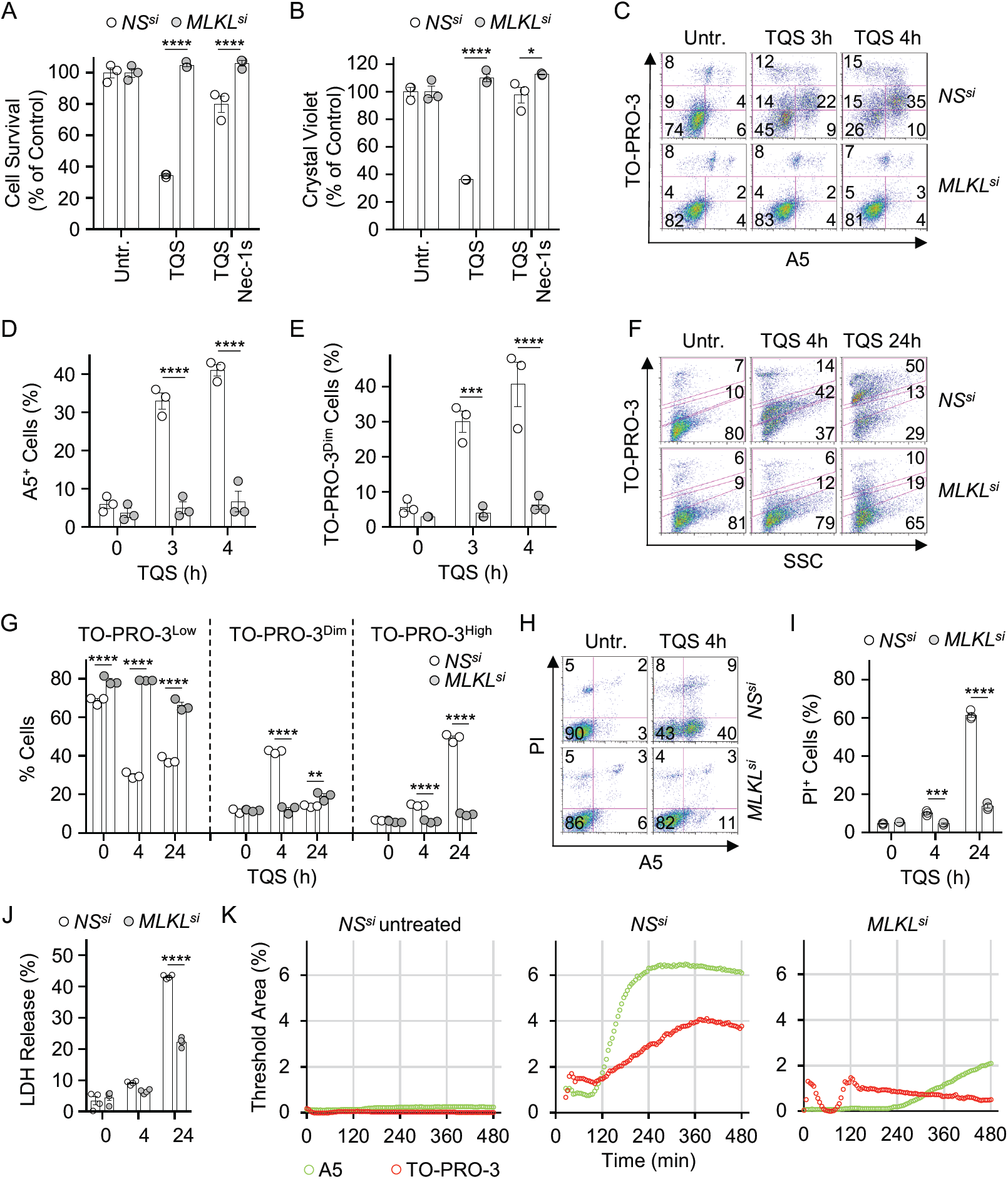
MLKL controls the selective Permeabilization of the Plasma Membrane during the early Stages of Necroptosis. HT-29 cells were transfected with siRNA for MLKL or scramble non-specific (NS) siRNA for 72 hours. Cells pre-treated with 10 μM QVD-OPh (Q), 5 μM Birinapant (S), 20 μM necrostatin-1 (Nec-1s) were stimulated with 1-10 ng.mL^−1^TNF*α* (T). **(A) and (B),** Cell viability was assessed by CellTiter-Glo (A), and Crystal Violet (B) after 16 hours. Data are means ± SEM of three independent experiments. \**P*< 0.1, \*\**P*<0.01, \*\*\*\**P*<0.0001 (ANOVA). **(C-E),** Flow cytometric analysis of cells stained with TO-PRO-3 and Annexin V (A5). Histograms show means ± SEM of three independent experiments. \*\*\**P*<0.001, \*\*\*\**P*<0.0001 (ANOVA). **(F-I),** Cells were collected after 4 and 24 hours of treatment, and stained with TO-PRO-3 and A5 (F and G), or with propidium iodide (PI) and A5 (H and I). \*\**P*<0.01, \*\*\**P*< 0.001, \*\*\*\**P*<0.0001 (ANOVA). (J), Measurement of lactate deshydrogenase (LDH) released from necroptotic cells, as indicated (means ± SEM, n=4, \*\*\*\**P*<0.0001, ANOVA). **(K),** Time lapsed microscopy analysis of A5 and TO-PRO-3 staining upon induction of necroptosis. The presented data are representative of at least three independent experiments.

### Pannexin-1 governs Plasma Membrane “Leakiness”

This “leakiness” of necroptotic cells is reminiscent of the caspase-mediated opening of PM PANX1 hemichannels during apoptosis, as evidence by the selective uptake of TO-PRO-3 [21]. This results in the release of purines, such as ATP in the extracellular space, which serve as a “find-me” signal for surrounding phagocytes [21]. To test the impact of PANX1 on “leakiness” during necroptosis, we first knocked down its expression with siRNA and found no overt impact on the phosphorylation of the necrosome core components RIPK1, RIPK3 and MLKL (Fig. 2A). Of note, we observed the appearance of a shorter MLKL fragment 4-6 hours post-stimulation, which was PANX1-dependent (Fig. 2A and 2B). The size of the generated fragment and the specificity of antibodies used (Fig. S1) suggested a cleavage between the brace region and the pseudokinase domain of MLKL [22] (see below). Nevertheless, oligomerization of MLKL occurred normally in PANX1-silenced cells, and was efficiently blocked by pre-treating cells with Nec-1s (Fig. 2B). In line with this, no significant change in the accumulation of phospho-MLKL at the PM was observed (Fig. 2C). Moreover, PANX1 silencing did not alter necroptotic death (Fig. 2D). Accordingly, MLKL-mediated flip of PS occurred normally without PANX1, reinforcing the idea that PANX1 is dispensable for cell death by necroptosis (Fig. 2E and 2F). In contrast, silencing PANX1 was as efficient as knocking down MLKL in preventing TO-PRO-3 uptake (Fig. 2E and 2F). Importantly, similar results were obtained in cells treated with the PANX1 pharmacologial inhibitor Trovafloxacin [23] (Fig. 2G-I). Collectively, these data demonstrate that MLKL initiates “leakiness” of the PM via PANX1, and that PANX1 activation is dispensable for the exposure of PS during necroptosis.

**Figure 2.**
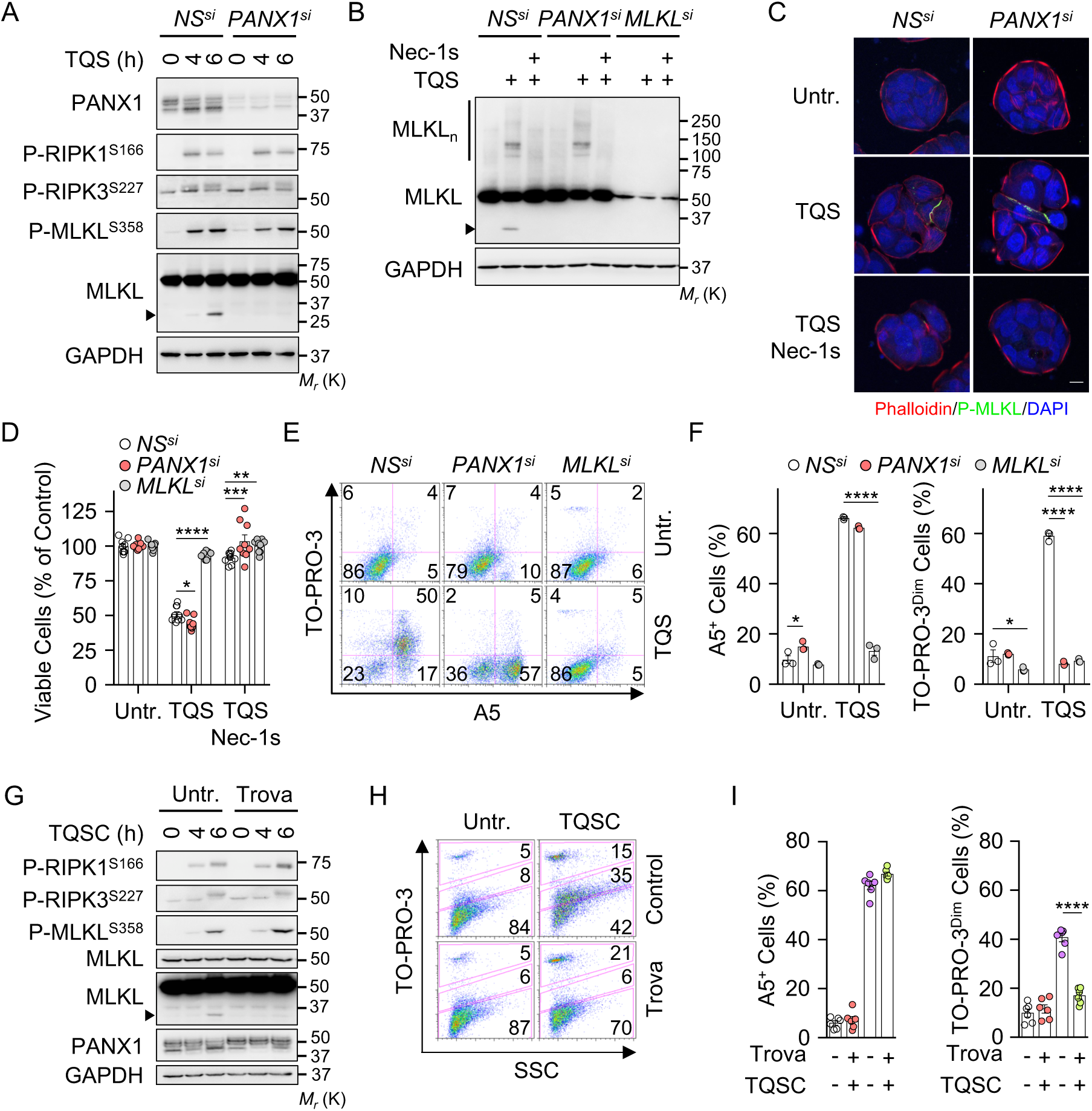
Pannexin-1 controls the Uptake of TO-PRO-3 during Necroptosis. HT-29 cells were transfected with siRNA for PANX1, MLKL, or scramble non-specific (NS) siRNA for 72 hrs. Cells were pre-treated with 10 μM QVD-OPh (Q) together with 5 μM Birinapant (S), prior stimulation with 10 ng.mL^−1^of TNF*α* (T), as indicated. Necrostatin-1 (Nec-1s, 20 μM) was also used. **(A) and (B),** Analysis of necroptosis hallmarks by Western blotting, as indicated. MLKL oligomers (MLKL_n_) were resolved by non-reducing SDS-PAGE after cross-linking. The arrowhead indicates MLKL cleaved fragment. On (B), cells were exposed to TQS for 5 hours. Molecular weight markers (M_r_) are shown. **(C),** confocal microscopic analysis of cells showing the localization of cytoskeletal actin with Phalloidin (red) and phosphorylated MLKL (green), with nuclei stained by DAPI (blue). Scale bar, 10 μm. **(D),** Cell viability assay by CellTiter-Glo in cells treated with TQS for 24 hours (mean ± SEM, n=9, \**P*<0.1, \*\**P*<0.01, \*\*\**P*<0.001, \*\*\*\**P*<0.0001, ANOVA). **(E) and (F),** Cells were treated with TQS for 4 hours, stained with TO-PRO-3 and Annexin V (A5), and analyzed by flow cytometry. Dead cells (TO-PRO-3^high^) were not analyzed. Data are means ± SEM of three independent experiments. \*\*\**P*<0.001, \*\*\*\**P*<0.0001 (ANOVA). **(G),** Western blot analysis of lysates from HT-29 cells treated as in (A) with the addition of 10 μM cycloheximide (C) to enhance cell death. When indicated, cells were pretreated with 30 μM of the PANX1 inhibitor Trovafloxacin (Trova). **(H) and (I)**, Flow cytometric analysis as indicated of cells treated as in (G) for 4 hours. Data are means ± SEM (n=6, \*\*\*\**P*<0.0001, ANOVA). The presented data are representative of at least three independent experiments.

During apoptosis, effector caspases remove a COOH-terminal inhibitory domain of PANX1 to irreversibly open the channels [21]. The analysis of PANX1 by immunoblotting revealed multiple species, consistent with *N*-glycosylation, as they were efficiently removed with a treatment with PNGase F ([24], Fig. 2A, and Fig. S2A). Although less dramatic than during apoptosis [21,25], PANX1 was also cleaved during necroptosis at a site close to the characterized caspase site, as evidenced by a slight decrease in full-length PANX1 combined with the presence of a cleaved fragment in PNGase F-treated samples (Fig. 2A, 2G and S2A). Conversely, the canonical caspase substrate PARP remained intact, arguing against residual caspase activity (Fig. S2A and S2B). Importantly, PANX1 proteolysis was significantly reduced in Trovafloxacin-treated cells during necroptosis but not apoptosis (Fig. 2G, S2B and S2C). Similarly, MLKL processing was hampered when PANX1 was silenced, or when its activity was blocked (Fig. 2A, 2B, and 2G). Hence, the cleavage of MLKL and PANX1 likely results from the opening of the channel in necroptotic cells.

### MLKL Oligomerization controls Plasma Membrane “Leakiness”

Once phosphorylated by RIPK3, MLKL binds high-order inositol phosphates [11]. This displaces MLKL’s auto-inhibitory domain and allows MLKL to oligomerize and reach the plasma membrane to exert its lethal function [11,12]. As expected [3,12], the treatment of cells with the small molecule necrosulfonamide (NSA) prevented MLKL oligomerization (Fig. 3A). NSA also significantly reduced the uptake of TO-PRO-3 (Fig. 3B and S3A). We next silenced inositol-tetrakisphosphate 1-kinase (ITPK1), a crucial kinase for the generation of inositol phosphates [11]. Although RIPK1, RIPK3, and MLKL were phosphorylated, the processing of MLKL and PANX1 no longer occurred without ITPK1 (Fig. 3C). In agreement with Dovey *et al* [11], silencing ITPK1 precluded MLKL oligomerization (Fig. 3D). This led to a reduction in the exposure of PS residues together with a significant inhibition of TO-PRO-3 uptake (Fig. 3E and 3F). Thus, MLKL oligomerization is instrumental for “leakiness” of the PM.

**Figure 3.**
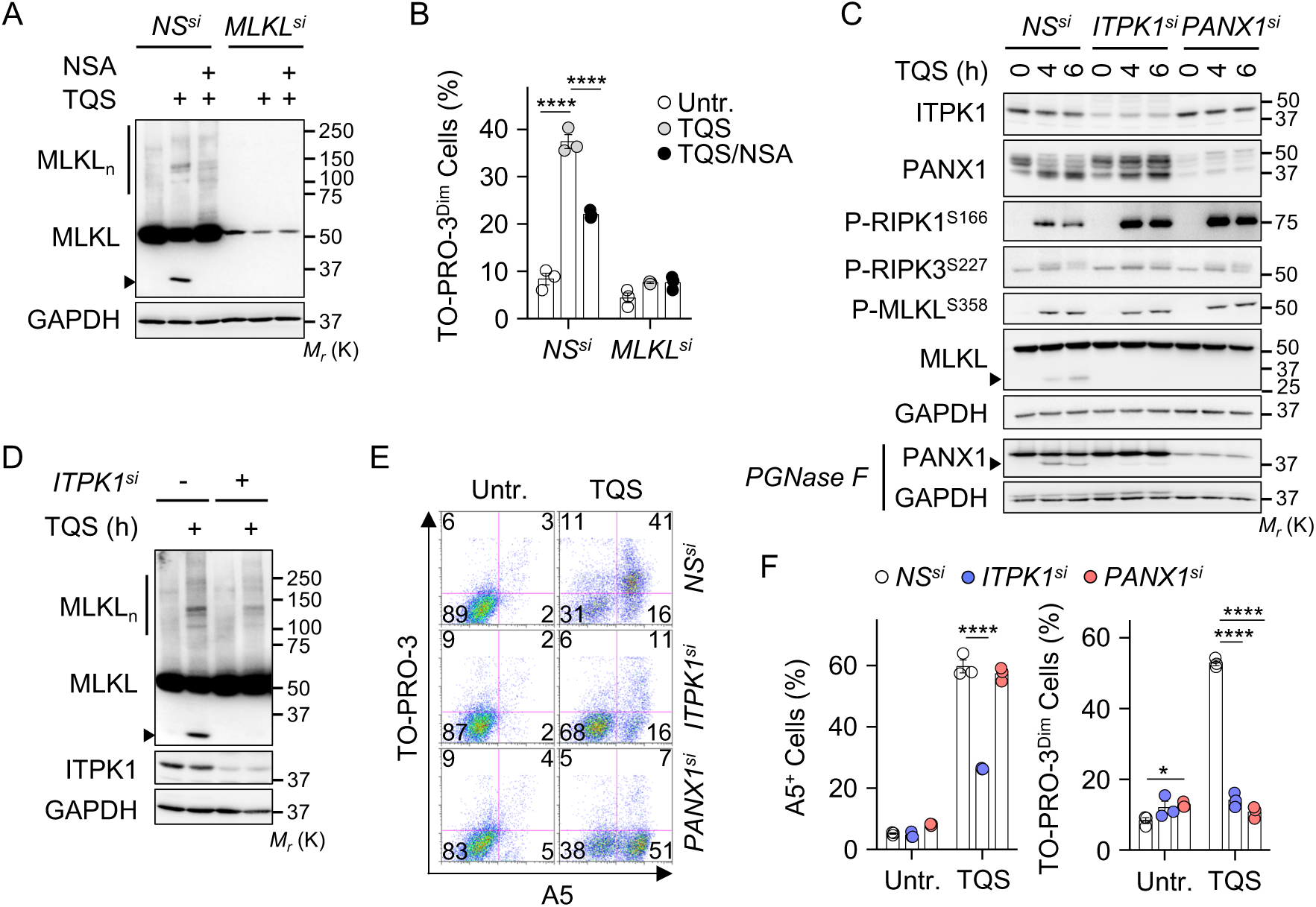
MLKL Oligomerization enables Plasma Membrane “Leakiness”. HT-29 cells were transfected with a nonspecific (NS) siRNA or with the indicated siRNAs for 72 hrs. Cells were pre-treated with 10 μM QVD-OPh (Q) plus 5 μM Birinapant (S), and exposed to 10 ng.mL^−1^of TNF*α* (T). **(A) and (B),** NS- and MLKL-silenced cells were treated with 5 μM necrosulfonamide (NSA) to inhibit MLKL oligomerization. (A) MLKL oligomerization was determined by non-reducing SDS-PAGE after cross-linking proteins. Cells were exposed to TQS for 5 hours. Arrowhead, MLKL cleaved fragment. Molecular weight markers (M_r_) are shown. (B) TO-PRO-3 uptake was analysed by flow cytometry in cells treated with TQS for 4 hours. Data are means ± SEM of three independent experiments. \*\*\*\**P*<0.0001 (ANOVA). **(C),** Cell lysates from NS-, ITPK1- and PANX1-silenced cells were analyzed by Western blotting with antibodies against the indicated targets. When indicated, samples were treated with PGNase F to remove glycosylation. **(D),** Western blotting analysis of MLKL oligomerization was performed as described in (A). **(E) and (F),** TO-PRO-3/Annexin V (A5) flow cytometric analysis of NS-, ITPK1-, and PANX1-silenced cells treated with TQS for 4 hours. Dead TO-PRO-3^high^cells were discarded. Data are means ± SEM of three independent experiments. \*\*\*\**P*<0.0001 (ANOVA). The presented data are representative of at least three independent experiments.

### Interplay between the small EVs Machinery and Pannexin-1 during Necroptosis

In addition to triggering cell death, active MLKL enables the generation of small EVs and their shedding from necroptotic cells [15,18]. To test whether the small EVs machinery contributes to “leakiness” during necroptosis, we focused on Rab27a and Rab27b, as this subfamily of Rab GTPases facilitates the docking and fusion to the PM of multivesicular bodies (MVBs), which are enriched with ILVs [26]. First, 100,000 × *g* (100k) sedimented membrane vesicles fractions, purified from culture media of Rab27a- or Rab27b-silenced cells, were analyzed by tunable resistive pulse sensing technology (Fig. 4A). Supporting previous works [15,18], the triggering of necroptosis increased small EVs release and this was markedly reduced when Rab27a or Rab27b were knocked down. Silencing Rab27a did not alter classical hallmarks of necroptosis, such as RIPK1 and MLKL phosphorylation or MLKL oligomerization, whereas PS exposure and overall cell death were slightly reduced (Fig. 4B-D, and Fig. S5). Yet, the proteolysis of MLKL and PANX1, as well as the uptake of TO-PRO-3 were drastically reduced without Rab27a (Fig. 4B-D). Because Rab27a is also instrumental for lysosomal biology [27], we assessed whether interfering with the lysosomes prevents MLKL processing. However, the inhibition of either the vacuolar H^+^ATPase with Bafilomycin A1 and concanamycin A, or of cathepsins had no gross effect on PANX1-mediated MLKL proteolysis (Fig. S4). Of note, the knockdown of Rab27b also prevented PM “leakiness” while sparing MLKL activation during necroptosis (Fig. 4E-G, and Fig. S5). Combined, this data suggests that oligomerized MLKL initiates PS asymmetry independently of the small EV machinery. However, the Rab27-dependent generation of small EVs appears crucial for the activation of PANX1 channels and “leakiness”.

**Figure 4.**
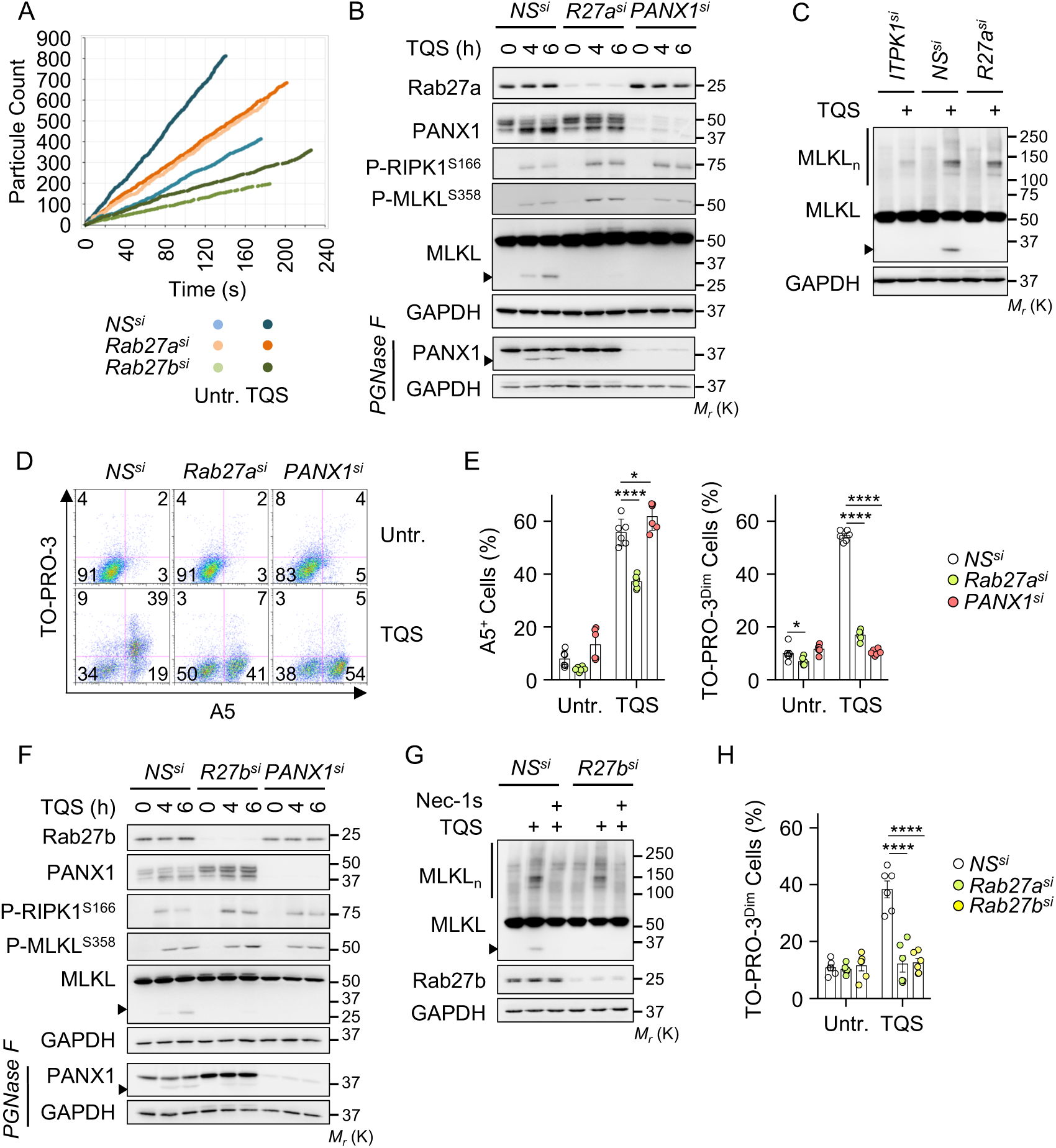
MLKL-mediated Pannexin-1 Activation requires the small EVs Machinery. **(A),** HT-29 cells were transfected with a nonspecific (NS) siRNA or with Rab27a- and Rab27b-specific siRNAs for 72 hrs. Cells were pre-treated with 10 μM QVD-OPh (Q) plus 5 μM Birinapant (S), and exposed to 10 ng.mL^−1^of TNF*α* (T) for 5 hours. Small EVs fractions were purified from the culture medium by ultracentrifugation. Shown is a representative diagram of particle count obtained with tunable resistive pulse sensing analysis (TRPS, qNano, IZON). **(B),** Cells as in (A) were subjected to Western blotting analysis with antibodies specific for the indicated proteins. When indicated, lysates were treated with PGNase F. The arrowhead shows cleavage products. Molecular weight markers (M_r_) are shown. **(C),** MLKL oligomerization was determined by non-reducing SDS-PAGE after protein cross-linking (TQS, 5 hours). **(D) and (E),** TO-PRO-3/Annexin V (A5) flow cytometric analysis of NS-, Rab27a-, and PANX1-silenced cells treated with TQS for 4 hours. Dead TO-PRO-3^high^cells were discarded. Data are means ± SEM (n=6, \**P*<0.1, \*\*\*\**P*<0.0001, ANOVA). **(F-H),** Cells were treated, and analyzed as in (B-E). The presented data are representative of at least three independent experiments.

### Pannexin-1 participates in the Intracellular Trafficking and the Secretion of small EVs

This connection between MVBs maturation and PANX1 prompted us to investigate the role of PANX1 in the release of small EVs. First, small EVs were purified from early necroptotic cells by ultracentrifugation (100k) and analyzed by immunoblot (Fig. 5A). As expected [15,18], necroptosis-induced small EVs release was MLKL-dependent, as evidence by staining the exosomal marker ALIX (Fig. 5A). In addition, PANX1, MLKL, and cleaved MLKL were also present in the 100k fraction upon necroptosis. Of note, the PANX1 band appeared slightly smaller, suggesting that it may be cleaved PANX1. Paralleling MLKL knockdown, silencing PANX1 also prevented the accumulation of the small EVs marker ALIX (Fig. 5B). Consistent with these biochemical analyses, the concentration of released vesicles was severely reduced without MLKL or PANX1, as measured by tunable resistive pulse sensing technology (Fig. 5C). Combined, these results suggest that PANX1 selectively controls the regulated secretion of small EVs during necroptosis.

**Figure 5.**
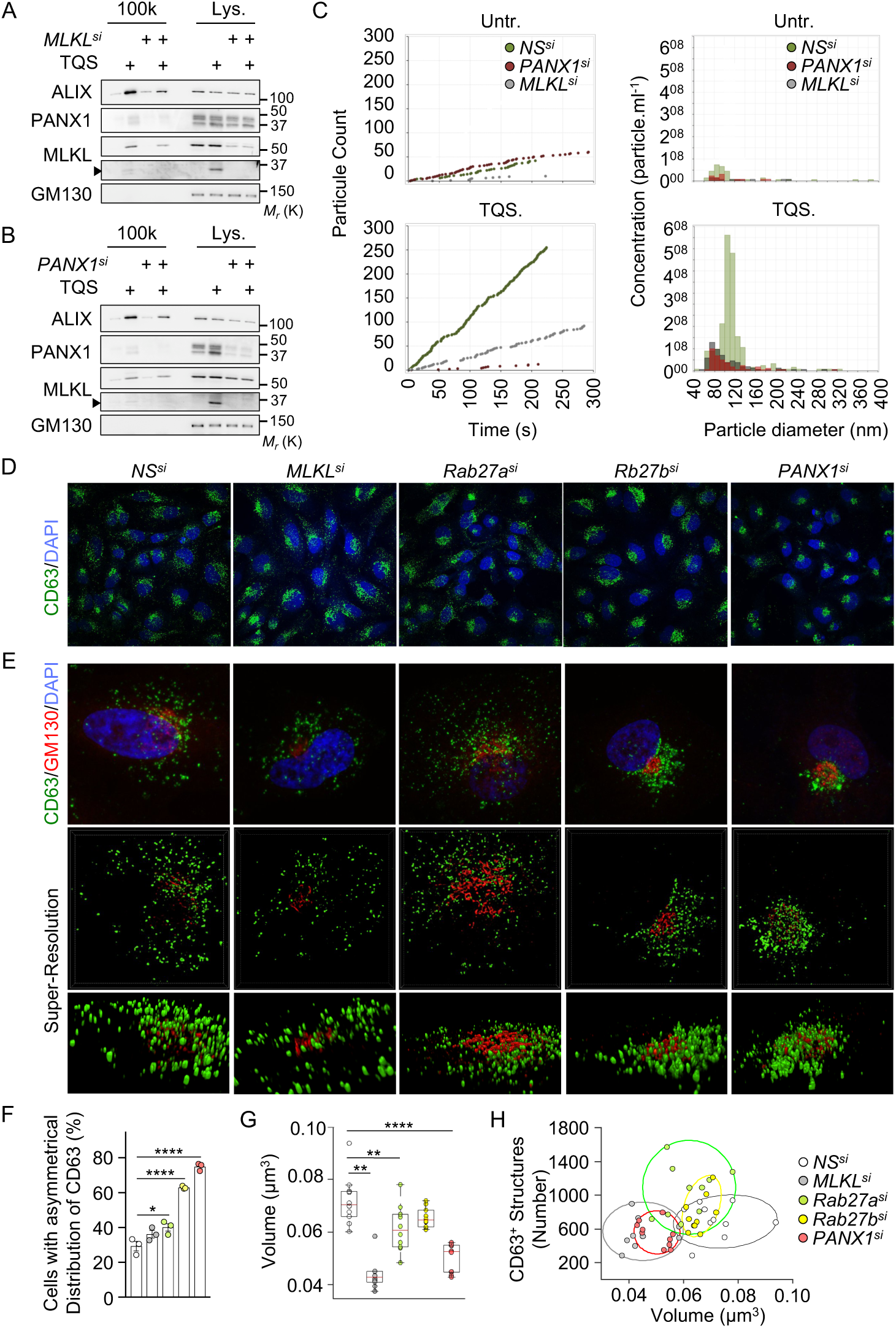
Pannexin-1 participates in the Generation and the Release of small EVs. HT-29 cells were transfected with a nonspecific (NS) siRNA or with the indicated siRNAs for 72 hrs. Cells were pre-treated with 10 μM QVD-OPh (Q) plus 5 μM Birinapant (S), and exposed to 10 ng.mL-1 of TNF*α* (T) for 6 hours. **(A and B,** small EVs were prepared and subjected to Western blotting analysis with antibodies specific for the indicated proteins. The Golgi resident protein GM130 served as a purity marker. Cleaved MLKL is indicated with an arrowhead, and molecular weight markers (M_r_) are shown. **C,** Diagram of size distribution in small EVs fractions from NS-, PANX1-, and MLKL-silenced cells, obtained with tunable resistive pulse sensing (TRPS, qNano, IZON). **D and E,** Confocal microscopic analysis of CD63 (D), or CD63 and GM130 (E) in HeLa cells, as indicated. In (E), staining was analyzed by Structure Illumination microscopy (SIM). Nuclei were counterstained with DAPI. **F,** The percentage of cells with an asymmetric distribution of CD63 was determined in three independent experiments. Data are means ± SEM (\**P*<0.1, \*\*\*\**P*<0.0001, ANOVA). **G and H,** Quantification of the volumes of CD63-positive structures (G), or scatter plots of numbers and mean volumes of CD63-positive structures analyzed by SIM (n=10 cells per condition; \*\**P*<0.05, \*\*\*\**P*<0.0001; ANOVA). Ellipses show 95% confidence. The presented data are representative of at least three independent experiments.

The absence of MLKL was shown to reduce the biogenesis of ILVs, and their constitutive release in the extracellular milieu [18]. The size and intracellular distribution of ILVs/MVBs is also significantly rearranged without Rab27 proteins [26]. To gain further insight into the mechanism responsible for the reduced small EVs release in PANX1-silenced cells, we first monitored, by confocal microscopy, the subcellular distribution of CD63, a tetraspanin enriched in ILVs [26]. This showed that PANX1-silenced cells phenocopied Rab27b knockdown and displayed an asymmetrical perinuclear accumulation of CD63-positive structures in HeLa cells (Fig. 5D-F, and Movies S3-S7). Of note, these CD63-positive vesicles clustered in closed proximity to the Golgi. The size and morphology of these structures were further analyzed by super-resolution microscopy. Supporting Ostrowski *et al* [26], Rab27a silencing, and not Rab27b knockdown, increased the number and the size heterogeneity of CD63-positive vesicles (Fig. 5G, 5H, and S6). By contrast, the number of vesicles dropped without MLKL, consistent with a defect in ILVs biogenesis [18]. Without PANX1, we observed a significant reduction in the number of CD63-positive structures, although their volume was not overtly changed (Fig. 5G, 5H, and S6). Altogether, these findings support the idea that PANX1, along with MLKL and the Rab27 proteins, contributes to the biogenesis and intracellular trafficking of small EVs.

## DISCUSSION

Here, we provide evidence that PANX1 channels operate downstream of MLKL to favor necroptosis-driven release of small EVs (Fig. S8). Our data also illustrate how the PM of necroptotic cells undergoes drastic rearrangements before it collapses. This includes the insertion of MLKL oligomers [10–14], PS scrambling, ions fluxes [13,28], activation of cell surface proteases [29], the release of small “bubbles” of broken PM [16], and PANX1-mediated uptake of dye and small EVs release. Although PANX1 silencing only marginally affected PS asymmetry, additional work is required to elucidate whether PS externalization is a prerequisite for PM “leakiness”, or if MLKL signaling bifurcates.

How exactly is MLKL linked to PANX1? MLKL oligomers target intracellular compartments, in addition to the PM [3]. Whether MLKL’s location dictates its apparently independent functions remains unknown. Nevertheless, our results indicate that MLKL oligomers-mediated activation of PANX1 hemichannels requires the small EVs machinery. Future investigations will define how precisely docking and fusion of MVBs to the PM is propitious for PM “leakiness”. In addition to mechanical stretches, ATP, K^+^efflux, Ca^2+^influx, phosphorylation, or caspase-mediated processing of its COOH-terminal intracellular latch were shown to open PANX1 channels [30]. Of note, oligomerized MLKL promotes a drastic efflux of potassium in bone marrow-derived macrophages, which culminates in the activation of the NLRP3 inflammasome [31]. However, buffering the concentration of K^+^with KCl at concentrations that did not interfere with MLKL phosphorylation (Ref. [31]) had no clear impact on PANX1-mediated uptake of TO-PRO-3 and processing of MLKL (Fig. S7). Although not formally ruled out here, MLKL-dependent Ca^2+^flux was shown to occur rather lately in necroptotic HT-29 cells, concomitantly to the loss of PM integrity [28]. Intriguingly, we noted that PANX1 and MLKL were cleaved by non-caspase proteases in a PANX1-dependent manner during necroptosis. Proteases contained in small EVs may be involved [32]. Interestingly, MLKL phosphorylation and PS asymmetry do not mark a point-of-no-return and both events can be reversed if the insult is removed [15–17]. Because “leakiness” occurs concomitantly to PS exposure, it is tempting to speculate that PANX1 opening during necroptosis is also reversible.

In addition to convey material for intercellular communication, necroptotic small EVs were proposed to counteract cell death by expelling lethal molecules such as active MLKL [18]. However, we find that curbing EVs release downstream of MLKL, by targeting Rab27a, Rab27b or PANX1 did not change the overall threshold for necroptotic death. Instead, we observed a slight reduction in cell death without Rab27 isoforms. During apoptosis, the presence of PS on the outer leaflet of the PM, combined with the opening of PANX1 hemichannels serve as “eat-me” and “find-me” signals to favor silent phagocytosis [21,23,33]. Caspase-mediated activation of PANX1 also coordinates the release of apoptotic bodies and prevents the formation of “string-like” protrusion called apoptopodia [23,34]. During necroptosis, the release in the extracellular milieu of “danger signals” from dying cells, together with ATP and small EVs drive inflammation [35]. Our data linking PANX1 to the biogenesis and release of small EVs suggest that tuning of PANX1 activity may shape the necroptotic microenvironment. The necroptotic machinery has been linked to several inflammatory conditions such as ischemia/reperfusion, inflammatory neurodegenerative diseases, intestinal inflammation, and cancer [1,36,37]. Strategies aimed at targeting PANX1 might be relevant to selectively modulate cell death-independent functions of MLKL.

## MATERIALS AND METHODS

### Cell Culture and Reagents

HT-29 and HeLa cells were purchased from the American Type Culture Collection. Bafilomycin A1, Concanamycin A, Cycloheximide, and Trovafloxacin were purchased from Sigma. Q-VD-OPh, Tumor necrosis factor-alpha (TNF*α*) were obtained from R&D Systems. Cathepsin Inhibitor 1 (S2847), Birinapant, Necrostatin-1 (Nec-1s), and Necrosulfonamide (NSA) were from Selleckchem. MCC-950 (Invivogen), TO-PRO-3, Alexa-488 Annexin V, and propidium iodide (Life Technologies), CellTiter-Glo and PGNase F (Promega), and Cytotoxicity Detection kit^PLUS^(Roche Diagnostics) were also used.

### siRNA, and Transfections

Cells were transfected with 10 pmol of siRNA using the Lipofectamine RNAiMAX Transfection Reagent (Invitrogen). The following sequences were obtained from Life Technologies: ITPK1, AGAUCCUGUCGUCUUCCAUGUAGGC (HSS105602); MLKL, GCAACGCAUGCCUGUUUCACCCAUA (HSS136796); PANX1, AUAUGAAUCCACAAAGGCAGCCUGA (HSS119236). A mixture of siRNAs targeting Rab27a was purchased from Sigma (EHU91501). Rab27b was silenced as previously published [38]: TAGGAATAGACTTTCGGGAAA.

### Cell Death Induction and Measurement

To induce necroptosis, HT-29 cells were pre-treated 30-60 min with 10 μM Q-VD-OPH plus 5 μM Birinapant prior stimulation with 0-10 ng.mL^−1^of TNF*α*. In some experiments, Nec-1s (20 μM), Trovafloxacin (30 μM), NSA (5 μM) and cycloheximide (10 μM) were used. Cell viability was assessed using CellTiter-Glo, staining with Crystal violet, and staining with Annexin V and TO-PRO-3, following the manufacturer instructions.

### Western Blotting Analysis and Antibodies

Cells were washed with ice-cold PBS and lysed with TNT buffer [50 mM Tris-HCl (pH 7.4), 150 mM NaCl, 1% Triton X-100, 1% Igepal, 2 mM EDTA] or with RIPA buffer [25 mM Tris-HCl (pH 7.4), 150 mM NaCl, 0.1% SDS, 0.5% Na-Deoxycholate, 1% NP40, 1 mM EDTA] supplemented with protease inhibitors (ThermoFisher Scientific) for 30 min on ice. Extracts were cleared by centrifugation at 9,000 × *g* and protein concentration determined by BCA (ThermoFisher Scientific). 5-10 μg proteins were resolved by SDS-polyacrylamide gel electrophoresis (SDS-PAGE) and transferred onto nitrocellulose membranes (GE Healthcare). To visualize MLKL oligomers, protein cross-linking was performed as previously described [39]. Briefly, cells were fixed in PBS containing 0.4% paraformaldehyde for 12 min at room temperature and lysed with PBS containing 1% Triton-X100 and proteases inhibitors. Proteins were incubated with 2X Laemmli buffer (Life Technologies) at room temperature and separated by SDS-PAGE.

Antibodies specific for the following proteins were purchased from Cell Signaling Technology: MLKL (D216N), P-MLKL S^358^ (D6H3V), PANX1 (D9M1C), Rab27a (D7V6B), P-RIPK1 S^166^ (D1L3S), RIPK3 (E1Z1D), and P-RIPK3 S^227^ (D6W2T). Antibodies specific for GAPDH (6C5), and PARP (F-2) were purchased from Santa Cruz Biotechnology. Antibodies specific for GM130 (ab52649), and for MLKL (ab183770) were purchased from Abcam. Antibodies against RIPK1 (BD Biosciences, 51), against MLKL (Millipore, MABC604), against ALIX (Biolegend, 3A9), against ITPK1 (Sigma, HPA055230), against CD63 (SBI, EXOAB A-1), and against Rab27b (Proteintech, 13412-1-AP) were also used

### Immunofluorescence Microscopy

HT-29 grown on coverslips were fixed with PBS containing 4% paraformaldehyde for 30 min at room temperature, and staining was performed as previously described [11]. Briefly, fixed samples were incubated at 4°C overnight with primary antibodies and 60 minutes with secondary antibodies in PBS containing 3% BSA, 1% saponin, and 1% Triton X-100). The following antibodies were used: anti-CD63 (556019, BD Biosciences), anti-P-MLKL S358/T357 (Abcam, ab187091), anti-GM130 (Abcam, ab52649). Alexa-568 Phalloidin (Life Technologies) was used to stain actin. Coverslip were sealed with Prolong gold anti-fade mouting media (Life Technologies) that allowed illumination of the nuclei by DAPI staining. For confocal microscopy, images were acquired with a Nikon A1R confocal microscope using the NIS-Element Software and processed using the FIJI Software. Structure illumination microscopy (SIM) images were acquired with a Nikon N-SIM microscope. Z-stacks of 0.12 μm were performed using a 100x oil-immersion lens with a 1.49 numerical aperture and reconstructed in 3D using the NIS-Element Software. Reconstructed images with a maximal NIS score of 8 were further analyzed. For live imaging, transfected HT-29 cells were pre-treated with Q-VD-OPh, Birinapant and incubated with TO-PRO-3 and Annexin V. After 30 minutes, TNF*α* was added and images were recorded every 5 minutes for 8 hrs. Acquisition and analysis were performed on a Leica Timelapse DMI-8 using the MetaMorph software (version 7.5) (MicroPicell, SFR François Bonamy, France).

### Image Analysis

Reconstructed images validated with the SIMcheck plugin 1.2 were analyzed with Fiji ImageJ2 v1.52i [40]. A binary mask was applied to select images with a modulation contrast-to-noise ratio (MCNR) > 8. Individual spots corresponding to single CD63-positive structure were isolated with the classic watershed function of the MorpholibJ plugin [41] and pixels intensity was measured. Only structures with the MCNR > 12 were retained (Fig. S8).

### Small Extracellular Vesicles Analysis

Small EVs were purified as previously described [42]. Briefly, to avoid contamination with extracellular vesicles (EVs) from the FBS, serum was centrifuged at 100,000 × *g* for 16 hrs on a Beckman Coulter ultracentrifuge using the SW-41 Ti rotor. EVs were purified from the conditional media of 0.4 × 10^6^ transfected HT-29 cells via successive centrifugations (300 × *g* 5 min, 2,000 × *g* 10 min, 10,000 × *g* 30 min, 100,000 × *g* 2h). Pellets were washed in PBS-0.22 μm filtered, re-centrifuged at 100,000 × *g* for 2 hours and resuspended in a small volume of PBS-0.22 μm filtered. 100,000 × *g* steps were performed on a Beckman Coulter Centrifuge using the MLA-130 rotor. Diameter and concentration of the particles were determined by Tunable Resistive Pulse Sensing (TRPS) using the qNano system (IZON). Particles within the range of 50-330 nm were detected using the pore “NP 100”. A stretch 46 mm was applied to the pores with an appropriate voltage (0.80Ω-1.00V) to reach a 120 nA baseline with noise < 10 pA. For Western Blot analysis, the following antibodies and concentrations were used: ALIX (Biolegend, 3A9; 1:1000), GM130 (ab52649; 1:2000), MLKL (ab183770; 1:1000) and PANX1 (D9M1C, 1:1000). We have submitted all relevant data of our experiments to the EV-TRACK knowledgebase (EV-TRACK ID: EV180073) [43].

### Statistical Analysis

Statistical analyses were performed using GraphPad Prism 7 software using one-way and two-way analysis (ANOVA).

## Supplementary Materials

Fig. S1. MLKL processing during necroptosis.

Fig. S2. PANX1 cleavage occurs secondarily to PANX1 activation. Fig. S3. Necrosulfonamide inhibition of TO-PRO-3 uptake.

Fig. S4. MLKL cleavage is not affected by lysosome inhibitors.

Fig. S5. Differential impact of PANX1, Rab27a, and Rab27b on cell survival.

Fig. S6. Effect of MLKL, PANX1, Rab27a, and Rab27b on CD63-positive structures.

Fig. S7. High concentrations of potassium chloride do not alter PANX1 activation and MLKL processing.

Fig. S8. Working model for MLKL-Rab27-PANX1 involvement in small EVs release.

**Figure S1.**
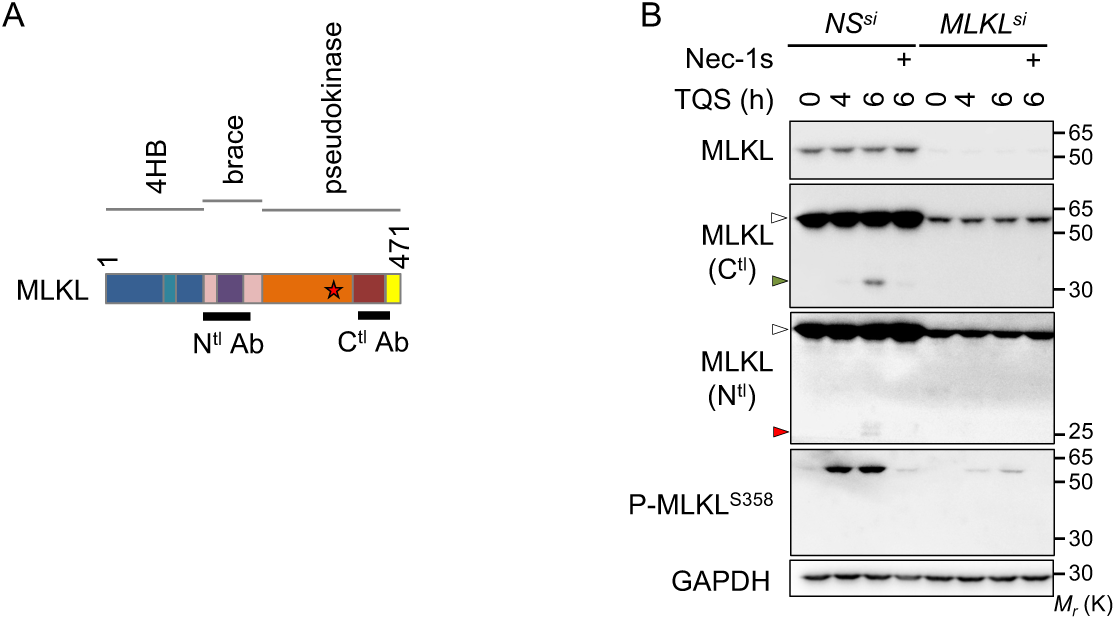
MLKL Processing during Necroptosis. **A,** Schematical representation of MLKL. The position of the epitopes recognized by the two antibodies used is shown. The star indicates the S358 position. **B,** HT-29 cells were transfected with siRNA for MLKL, or scramble non-specific (NS) siRNA for 72 hrs. Cells were pre-treated with 10 μM QVD-OPh (Q) together with 5 μM Birinapant (S), prior stimulation with 10 ng.mL^−1^of TNF*α* (T), as indicated. Necrostatin-1 (Nec-1s, 20 μM) was also used. Cell lysates were prepared and analyzed by immunoblot, as indicated. Molecular weight markers (Mr) are shown.

**Figure S2.**
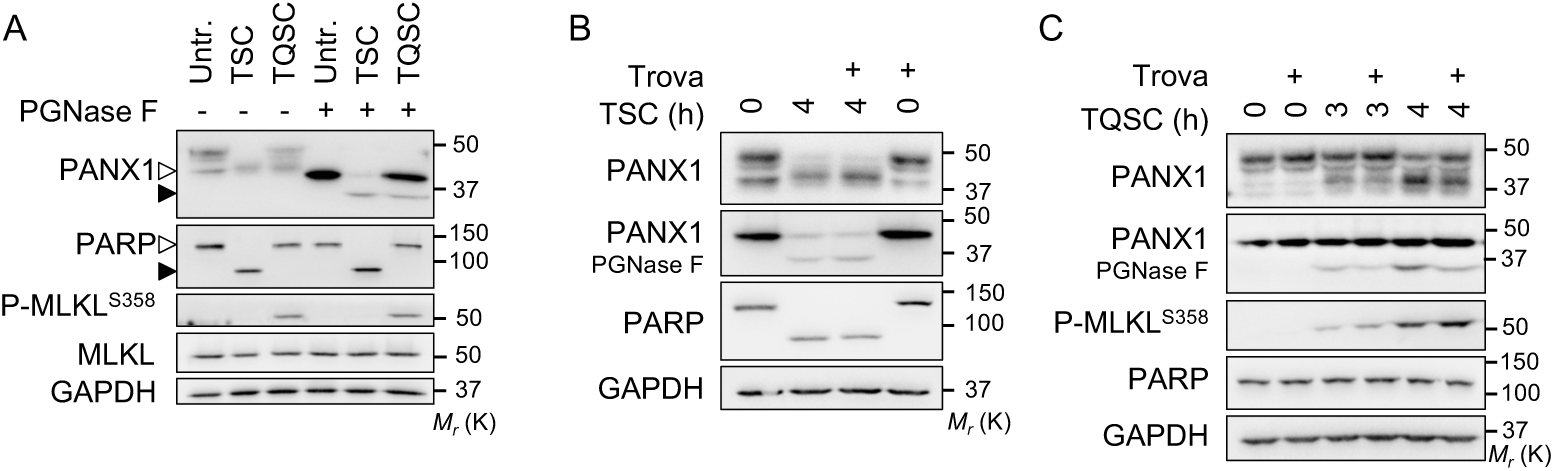
PANX1 Cleavage occurs Secondarily to PANX1 Activation. **A-C,** Western blot analysis of lysates from HT-29 cells treated with 10 μM QVD-OPh (Q), 5 μM Birinapant (S), 10 μM cycloheximide (C), prior stimulation with 10 ng.mL^−1^of TNF*α* (T), as indicated. When indicated, cells were pretreated with 30 μM of the PANX1 inhibitor Trovafloxacin (Trova). A treatment with PGNase F was used to remove glycosylation. Open and close symbols represent full-length and cleaved proteins, respectively. Molecular weight markers (Mr) are shown.

**Figure S3.**
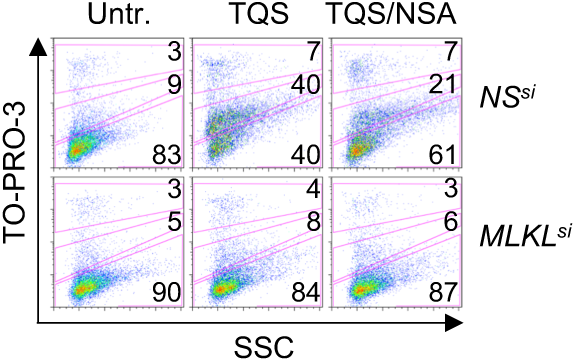
Necrosulfonamide Inhibition of TO-PRO-3 Uptake. Flow cytometric analysis, relative to Figure 3B.

**Figure S4.**
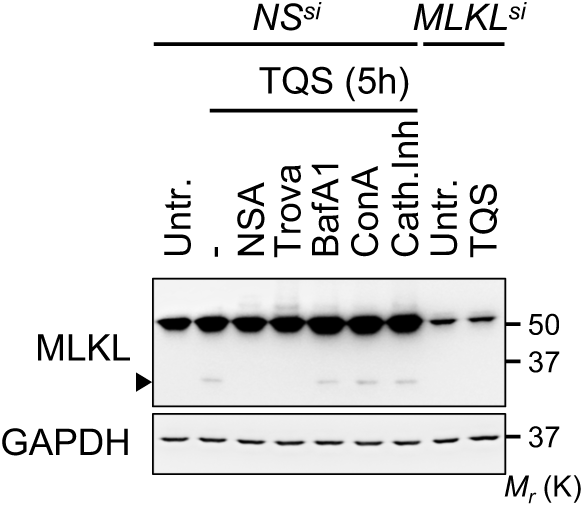
MLKL Cleavage is not affected by Lysosome Inhibitors. Control non-specific (NS) or MLKL-silenced HT-29 were pre-treated with 5 μM NSA, 30 μM Trovafloxacin (Trova), 100 ng.mL^−1^Bafilomycin A1, 10 nM concanamycin A1, 10 μM Cathepsin inhibitor 1, and necroptosis was triggered, as in Figure 1. MLKL processing was analysed by Western Blotting, as indicated. Molecular weight markers (Mr) are shown.

**Figure S5.**
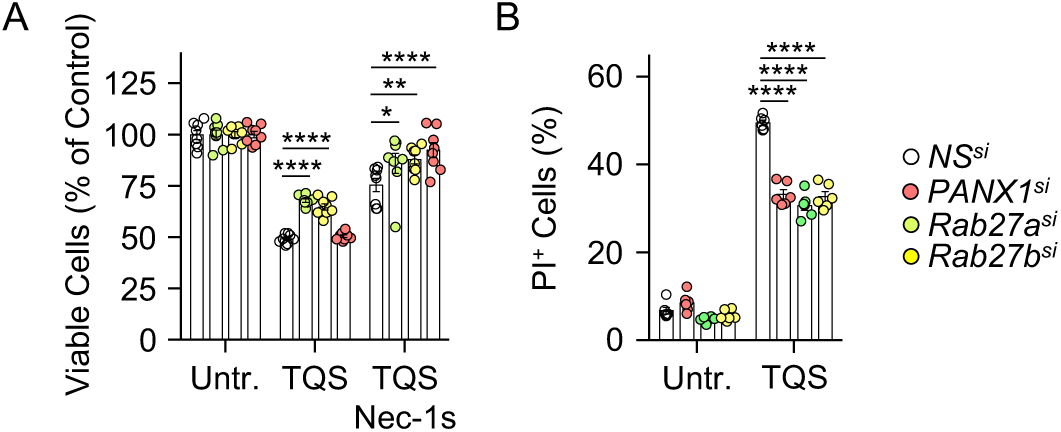
Differential Impact of PANX1, Rab27a, and Rab27b on Cell Survival. HT-29 cells were transfected with siRNA for PANX1, Rab27a, Rab27b, or scramble non-specific (NS) siRNA for 72 hrs. Cells were pre-treated with 10 μM QVD-Oph (Q) together with 5 μM Birinapant (S), prior stimulation with 10 ng.mL^−1^ of TNF*α* (T) for 24 h. Necrostatin-1 (Nec-1s, 20 μM) was also used. **A,** Cell viability assay by CellTiter-Glo (mean ± SEM, n=8, \**P*<0.1, \*\**P*<0.01, \*\*\*\**P*<0.0001, ANOVA). **B,** Flow cytometric analysis of cells stained with Propidium Iodide (PI). Shown is mean ± SEM (n=6, \*\*\*\**P*<0.0001, ANOVA).

**Figure S6.**
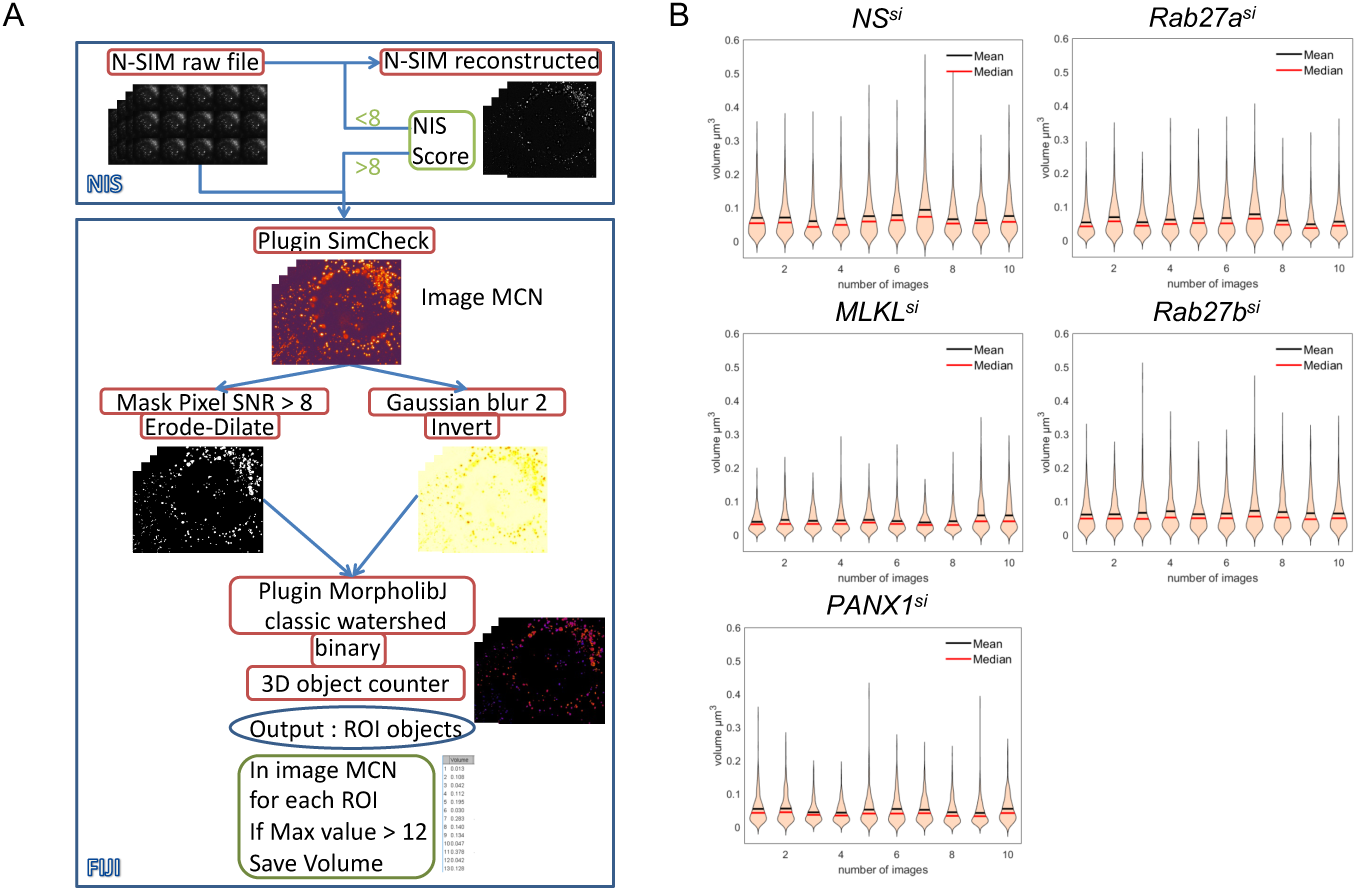
Effect of MLKL, PANX1, Rab27a, and Rab27b on CD63-positive Structures. **A,** Images analysis strategy. **B,** Density plots showing dispersion of the volume of CD63-positive structures for each cell.

**Figure S7.**
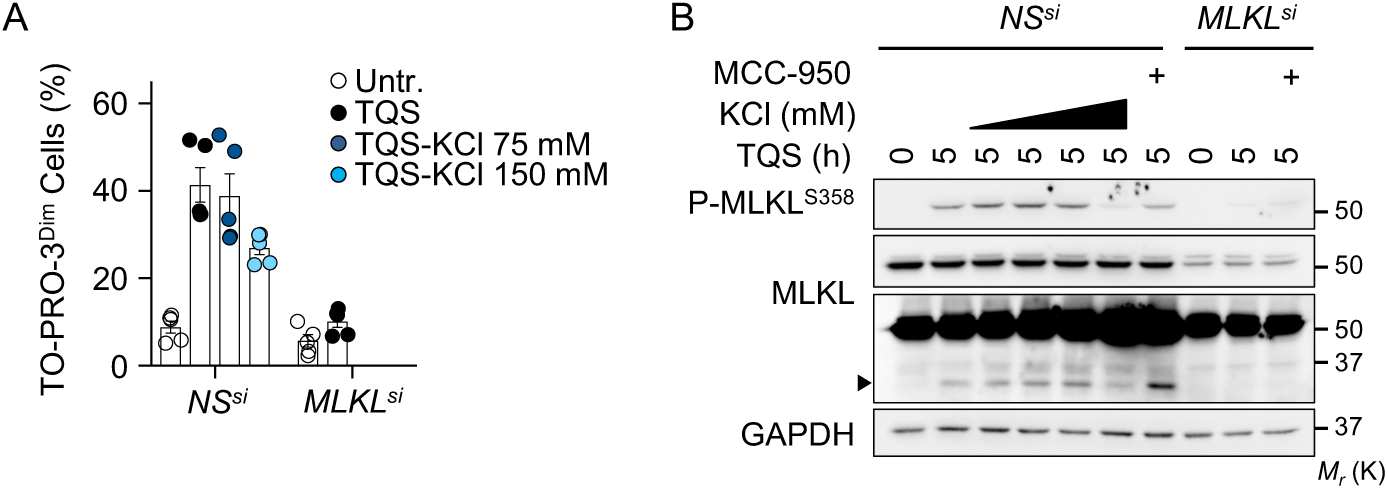
High Concentrations of Potassium Chloride do not alter PANX1 Activation and MLKL Processing. **A,** HT-29 cells were transfected with siRNA for MLKL, or scramble non-specific (NS) siRNA for 72 hrs. Cells were pre-treated with 10 μM QVD-OPh (Q) together with 5 μM Birinapant (S) and KCl, prior stimulation with 10 ng.mL^−1^of TNF*α* (T) for 4 h. Flow cytometric analysis of cells stained with TO-PRO-3 (mean ± SEM, n=5). **B,** Immunoblots, as indicated, of cells as in (A). MCC-950 (1 μM) was also used. Molecular weight markers (Mr) are shown.

**Figure S8.**
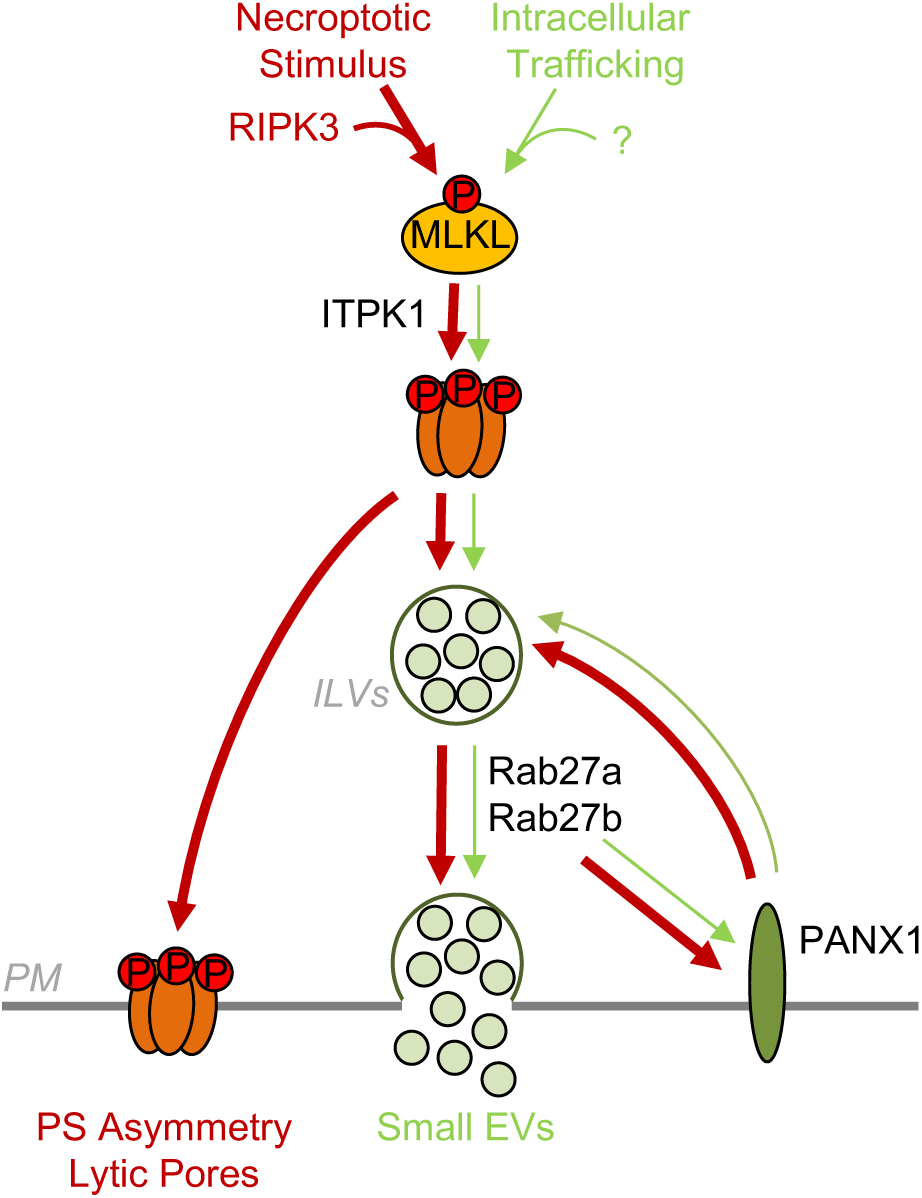
Working model for MLKL-Rab27-PANX1 involvement in small EVs Generation and Release. In viable cells, a portion of MLKL is constitutively phosphorylated and oligomerized, possibly in intracellular membranes. This drives the basal release of small EVs via the Rab27 subfamily and PANX1 hemichannels. During necroptosis RIPK3-mediated phosphorylation of MLKL increases this secretion pathway. Simultaneously, MLKL oligomerizes at the plasma membrane (PM) to promote phosphatidylserine (PS) asymmetry and form lytic pores.

Movies S1-2. MLKL prevents Annexin-V asymmetry and TO-PRO-3 uptake during necroptosis.

Movies S3-7. Differential impact of MLKL, PANX1, Rab27a, and Rab27b on CD63 staining.

## Acknowledgements

We thank Kathryn Jacobs for critically reading the manuscript; Micropicell and Cytocell facilities (SFR Santé François Bonamy, Nantes, France). This research was funded by an International Program for Scientific Cooperation (PICS, CNRS), Fondation pour la Recherche Medicale (Equipe labellisée DEQ20180339184), Fondation ARC contre le Cancer (NB), Ligue nationale contre le cancer comités de Loire-Atlantique, Maine et Loire, Vendée (JG, NB), Région Pays de la Loire et Nantes Métropole under Connect Talent Grant (JG). TD is a PhD fellow funded by Nantes Métropole; GAG holds postdoctoral fellowships from Fondation de France. The authors declare no competing interests.

## Author contributions

TD, GAG, MF, PH, JG, NB, conception and design, acquisition of data, analysis and interpretation of data, drafting or revising the article. All authors approved the manuscript.

## References

1. Shan B, Pan H, Najafov A, Yuan J (2018) Necroptosis in development and diseases. Genes Dev 32: 327–340.

2. Weinlich R, Oberst A, Beere HM, Green DR (2017) Necroptosis in development, inflammation and disease. Nat Rev Mol Cell Biol 18: 127–136.

3. Sun L, Wang H, Wang Z, He S, Chen S, Liao D, Wang L, Yan J, Liu W, Lei X, et al. (2012) Mixed lineage kinase domain-like protein mediates necrosis signaling downstream of RIP3 kinase. Cell 148: 213–227.

4. He S, Wang L, Miao L, Wang T, Du F, Zhao L, Wang X (2009) Receptor interacting protein kinase-3 determines cellular necrotic response to. Cell 137: 1100–1111.

5. Zhang D-W, Shao J, Lin J, Zhang N, Lu B-J, Lin S-C, Dong M-Q, Han J (2009) RIP3, an energy metabolism regulator that switches TNF-induced cell death from apoptosis to necrosis. Science 325: 332–336.

6. Degterev A, Huang Z, Boyce M, Li Y, Jagtap P, Mizushima N, Cuny GD, Mitchison TJ, Moskowitz MA, Yuan J (2005) Chemical inhibitor of nonapoptotic cell death with therapeutic potential for ischemic brain injury. Nat Chem Biol 1: 112–119.

7. Cho YS, Challa S, Moquin D, Genga R, Ray TD, Guildford M, Chan FK-M (2009) Phosphorylation-driven assembly of the RIP1-RIP3 complex regulates programmed necrosis and virus-induced inflammation. Cell 137: 1112–1123.

8. Zhao J, Jitkaew S, Cai Z, Choksi S, Li Q, Luo J, Liu Z-G (2012) Mixed lineage kinase domain-like is a key receptor interacting protein 3 downstream component of TNF-induced necrosis. Proc Natl Acad Sci U S A 109: 5322–5327.

9. Mompean M, Li W, Li J, Laage S, Siemer AB, Bozkurt G, Wu H, McDermott AE (2018) The Structure of the Necrosome RIPK1-RIPK3 Core, a Human Hetero-Amyloid Signaling Complex. Cell 173: 1244–1253.e10.

10. Murphy JM, Czabotar PE, Hildebrand JM, Lucet IS, Zhang J-G, Alvarez-Diaz S, Lewis R, Lalaoui N, Metcalf D, Webb AI, et al. (2013) The pseudokinase MLKL mediates necroptosis via a molecular switch mechanism. Immunity 39: 443–453.

11. Dovey CM, Diep J, Clarke BP, Hale AT, McNamara DE, Guo H, Brown NWJ, Cao JY, Grace CR, Gough PJ, et al. (2018) MLKL Requires the Inositol Phosphate Code to Execute Necroptosis. Mol Cell 70: 936–948.e7.

12. Wang H, Sun L, Su L, Rizo J, Liu L, Wang L-F, Wang F-S, Wang X (2014) Mixed lineage kinase domain-like protein MLKL causes necrotic membrane disruption upon phosphorylation by RIP3. Mol Cell 54: 133–146.

13. Cai Z, Jitkaew S, Zhao J, Chiang H-C, Choksi S, Liu J, Ward Y, Wu L-G, Liu Z-G (2014) Plasma membrane translocation of trimerized MLKL protein is required for. Nat Cell Biol 16: 55–65.

14. Dondelinger Y, Declercq W, Montessuit S, Roelandt R, Goncalves A, Bruggeman I, Hulpiau P, Weber K, Sehon CA, Marquis RW, et al. (2014) MLKL compromises plasma membrane integrity by binding to phosphatidylinositol phosphates. Cell Rep 7: 971–981.

15. Zargarian S, Shlomovitz I, Erlich Z, Hourizadeh A, Ofir-Birin Y, Croker BA, Regev-Rudzki N, Edry-Botzer L, Gerlic M (2017) Phosphatidylserine externalization, ‘necroptotic bodies’ release, and phagocytosis during necroptosis. PLoS Biol 15: e2002711.

16. Gong Y-N, Guy C, Olauson H, Becker JU, Yang M, Fitzgerald P, Linkermann A, Green DR (2017) ESCRT-III Acts Downstream of MLKL to Regulate Necroptotic Cell Death and Its Consequences. Cell 169: 286–300.e16.

17. Gong Y-N, Guy C, Crawford JC, Green DR (2017) Biological events and molecular signaling following MLKL activation during necroptosis. Cell Cycle 16: 1748–1760.

18. Yoon S, Kovalenko A, Bogdanov K, Wallach D (2017) MLKL, the Protein that Mediates Necroptosis, Also Regulates Endosomal Trafficking and Extracellular Vesicle Generation. Immunity 47: 51–65.e7.

19. Colombo M, Raposo G, Thery C (2014) Biogenesis, secretion, and intercellular interactions of exosomes and other extracellular vesicles. Annu Rev Cell Dev Biol 30: 255–289.

20. Galluzzi L, Kepp O, Chan FK-M, Kroemer G (2017) Necroptosis: Mechanisms and Relevance to Disease. Annu Rev Pathol 12: 103–130.

21. Chekeni FB, Elliott MR, Sandilos JK, Walk SF, Kinchen JM, Lazarowski ER, Armstrong AJ, Penuela S, Laird DW, Salvesen GS, et al. (2010) Pannexin 1 channels mediate ‘find-me’ signal release and membrane permeability during apoptosis. Nature 467: 863–867.

22. Hildebrand JM, Tanzer MC, Lucet IS, Young SN, Spall SK, Sharma P, Pierotti C, Garnier J-M, Dobson RCJ, Webb AI, et al. (2014) Activation of the pseudokinase MLKL unleashes the four-helix bundle domain to induce membrane localization and necroptotic cell death. Proc Natl Acad Sci U S A 111: 15072–15077.

23. Poon IKH, Chiu Y-H, Armstrong AJ, Kinchen JM, Juncadella IJ, Bayliss DA, Ravichandran KS (2014) Unexpected link between an antibiotic, pannexin channels and apoptosis. Nature 507: 329–334.

24. Boassa D, Ambrosi C, Qiu F, Dahl G, Gaietta G, Sosinsky G (2007) Pannexin1 channels contain a glycosylation site that targets the hexamer to the plasma membrane. J Biol Chem 282: 31733–31743.

25. Boyd-Tressler AM, Lane GS, Dubyak GR (2017) Up-regulated Ectonucleotidases in Fas-Associated Death Domain Protein-and Receptor-Interacting Protein Kinase 1-Deficient Jurkat Leukemia Cells Counteract Extracellular ATP/AMP Accumulation via Pannexin-1 Channels during Chemotherapeutic Drug-Induced Apoptosis. Mol Pharmacol 92: 30–47.

26. Ostrowski M, Carmo NB, Krumeich S, Fanget I, Raposo G, Savina A, Moita CF, Schauer K, Hume AN, Freitas RP, et al. (2010) Rab27a and Rab27b control different steps of the exosome secretion pathway. Nat Cell Biol 12: 19–30; sup pp 1-13.

27. Luzio JP, Hackmann Y, Dieckmann NMG, Griffiths GM (2014) The biogenesis of lysosomes and lysosome-related organelles. Cold Spring Harb Perspect Biol 6: a016840.

28. Ros U, Pena-Blanco A, Hanggi K, Kunzendorf U, Krautwald S, Wong WW-L, Garcia-Saez AJ (2017) Necroptosis Execution Is Mediated by Plasma Membrane Nanopores Independent of Calcium. Cell Rep 19: 175–187.

29. Cai Z, Zhang A, Choksi S, Li W, Li T, Zhang X-M, Liu Z-G (2016) Activation of cell-surface proteases promotes necroptosis, inflammation and cell migration. Cell Res 26: 886–900.

30. Chiu Y-H, Schappe MS, Desai BN, Bayliss DA (2018) Revisiting multimodal activation and channel properties of Pannexin 1. J Gen Physiol 150: 19–39.

31. Conos SA, Chen KW, De Nardo D, Hara H, Whitehead L, Nunez G, Masters SL, Murphy JM, Schroder K, Vaux DL, et al. (2017) Active MLKL triggers the NLRP3 inflammasome in a cell-intrinsic manner. Proc Natl Acad Sci U S A 114: E961–E969.

32. Shimoda M, Khokha R (2013) Proteolytic factors in exosomes. Proteomics 13: 1624–1636.

33. Segawa K, Nagata S (2015) An Apoptotic ‘Eat Me’ Signal: Phosphatidylserine Exposure. Trends Cell Biol 25: 639–650.

34. Atkin-Smith GK, Tixeira R, Paone S, Mathivanan S, Collins C, Liem M, Goodall KJ, Ravichandran KS, Hulett MD, Poon IKH (2015) A novel mechanism of generating extracellular vesicles during apoptosis via a beads-on-a-string membrane structure. Nat Commun 6: 7439.

35. Kearney CJ, Martin SJ (2017) An Inflammatory Perspective on Necroptosis. Mol Cell 65: 965–973.

36. Seehawer M, Heinzmann F, D’Artista L, Harbig J, Roux P-F, Hoenicke L, Dang H, Klotz S, Robinson L, Dore G, et al. (2018) Necroptosis microenvironment directs lineage commitment in liver cancer. Nature 562: 69–75.

37. Strilic B, Yang L, Albarran-Juarez J, Wachsmuth L, Han K, Muller UC, Pasparakis M, Offermanns S (2016) Tumour-cell-induced endothelial cell necroptosis via death receptor 6 promotes metastasis. Nature 536: 215–218.

38. Hendrix A, Maynard D, Pauwels P, Braems G, Denys H, Van den Broecke R, Lambert J, Van Belle S, Cocquyt V, Gespach C, et al. (2010) Effect of the secretory small GTPase Rab27B on breast cancer growth, invasion, and metastasis. J Natl Cancer Inst 102: 866–880.

39. Chiu Y-H, Jin X, Medina CB, Leonhardt SA, Kiessling V, Bennett BC, Shu S, Tamm LK, Yeager M, Ravichandran KS, et al. (2017) A quantized mechanism for activation of pannexin channels. Nat Commun 8: 14324.

40. Ball G, Demmerle J, Kaufmann R, Davis I, Dobbie IM, Schermelleh L (2015) SIMcheck: a Toolbox for Successful Super-resolution Structured Illumination Microscopy. Sci Rep 5: 15915.

41. Legland D, Arganda-Carreras I, Andrey P (2016) MorphoLibJ: integrated library and plugins for mathematical morphology with ImageJ. Bioinformatics 32: 3532–3534.

42. Andre-Gregoire G, Bidere N, Gavard J (2018) Temozolomide affects Extracellular Vesicles Released by Glioblastoma Cells. Biochimie.

43. Van Deun J, Mestdagh P, Agostinis P, Akay O, Anand S, Anckaert J, Martinez ZA, Baetens T, Beghein E, Bertier L, et al. (2017) EV-TRACK: transparent reporting and centralizing knowledge in extracellular vesicle research. Nat Methods 14: 228–232.

